# Prefrontal mechanisms of goal progress inference and monitoring in macaque monkeys

**DOI:** 10.64898/2026.05.19.726190

**Authors:** Xiaoqi Xu, Frederic M Stoll, Matteo di Volo, Charles R. E. Wilson, Emmanuel Procyk, Nils Kolling

## Abstract

Primates pursue goals over extended periods and over multiple steps. The ability to track progress and its rate is critical for such pursuit, yet the neural mechanisms of this tracking are unclear. We recorded single-unit activity from macaque midcingulate cortex (MCC) and lateral prefrontal cortex (LPFC) whilst animals worked for immediate rewards and checked a gauge showing progress towards a larger reward. MCC expressed a temporally extended progress rate signal linked to the long timescale neurons in the region. LPFC expressed progress rate only when inferences on that property were required. In MCC, independent feedback-valence information was also preserved, enabling both readout of work success and rate-specific progress updating. An RNN trained under the same constraints showed that progress rate representations emerge spontaneously, and are critical for tracking goal progress. These results demonstrate a frontal mechanism for inferring and predicting goal progress by extracting progress rate information.

## Introduction

Humans and other animals routinely pursue temporally extended goals such as completing a doctoral degree. Efficient long-horizon behaviour requires monitoring internal milestones (e.g., papers written, courses completed) while respecting external constraints (e.g., institutional requirements, unexpected results, deadlines). Progress monitoring cannot be reduced to evaluating recent outcomes - it also requires estimating how current actions translate into future progress, especially when environmental contingencies change. This ability is also a hallmark of primate behaviour, impaired across conditions, and closely associated with frontal cortex function.

A growing literature has identified neural correlates of progress-related variables in frontal and cingulate cortex. Medial frontal regions, including the midcingulate cortex (MCC), encode signals related to performance monitoring, behavioural adjustment, and persistence during extended actions^1–4^. These signals often reflect cumulative variables such as accumulated reward, effort expenditure, or proximity to a goal, consistent with a role in monitoring ongoing progress^5–7^. Complementarily, ventromedial prefrontal cortex (vmPFC) supports sustained goal commitment by selectively prioritizing goal-relevant information over competing alternatives, whereas lateral prefrontal cortex (LPFC) represents task structure, integrates information over time, and infers latent variables not directly observable from immediate feedback^9–12^.

However, most existing work operationalizes “progress” using proxies such as reward accumulation or elapsed time, leaving unresolved how the brain tracks progress rate itself: how quickly actions translate into advancement toward a distant goal. In natural environments, the action-to-progress mapping is rarely stationary. The same effort can yield different progress rates depending on external conditions (e.g., shifting task difficulty or environmental volatility). Adaptive control depends on inferring latent rate variables, including volatility, learning rates, and hazard rates, which govern how evidence, effort, or outcomes are integrated across time^13–15^. Yet whether and how the brain infers and represents a latent progress rate during extended goal-directed behaviour remains largely unknown.

We examined the neural mechanism of inferring and monitoring latent progress rates by analyzing single-unit recordings from MCC and LPFC in two macaques performing a task with variable, latent rates of incremental progress: the “check versus work” task^7^ (Fig. 1a). On each trial, animals choose either to work, performing a perceptual decision task for a small juice reward, or to check, revealing an on-screen gauge whose size reflects accumulated correct work trials. When the gauge reaches its maximum, a large bonus becomes available if the monkey checks again. Critically, the rate at which the gauge grows varies unpredictably across blocks. Thus, the speed of advancement toward the bonus is not directly observable and must be inferred through checking. Progress rate is therefore a latent contextual variable that shapes optimal decisions about when to work versus check.

**Figure 1.**
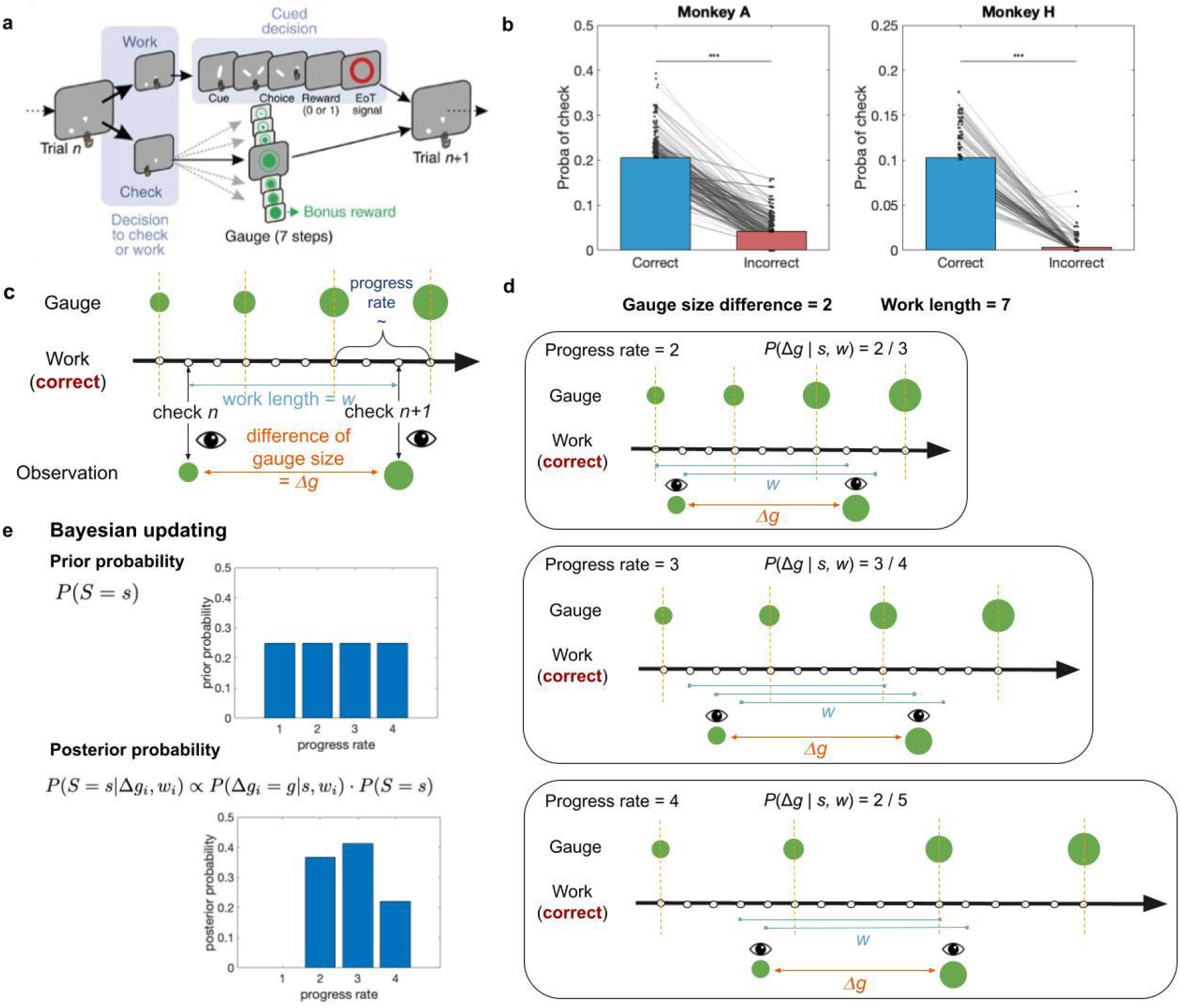
Task design and Bayesian inference of latent progress rate. **(a)** *Task structure (“check vs work”)*. On each trial, the monkey chooses to work or check. Work trials require a cued delayed perceptual decision and yield a small juice reward if correct. Check trials reveal the current gauge (7 discrete steps) and, when the gauge is full, deliver a bonus reward. Gauge growth depends on correct work and occurs at a block-specific, uncued progress rate. **(b)** *Behavioural contingency linking correct work to checking*. Probability of choosing check on the next opportunity as a function of the immediately preceding work outcome (correct vs incorrect), shown separately for each monkey. Points/lines indicate session-level values; bars show mean across sessions. Both monkeys check far less after incorrect outcomes, consistent with understanding that progress depends on correct work. **(c)** *Progress-rate manipulation and observable evidence between checks*. Progress rate is defined as the number of correct work trials required per one-unit gauge increase. Between two consecutive checks *(check n and check n+1), the animal can combine the work length w (number of correct work trials between checks; blue arrow)* with the observed gauge increment *Δg* (difference in observed gauge size between checks; orange) to infer the latent progress rate. **(d)** *Likelihood under a discrete gauge-growth process*. Example for *w=7* and *Δg=2*. For candidate progress rates *s*, the same observation can arise from multiple hidden alignments of the within-step counter at check time. Enumerating these alignments (assuming checks are uniformly distributed over possible counter phases) yields P(*Δg*|*s,w*) for each candidate rate (illustrated for *s=2,3,4*). **(e)** *Bayesian updating*. A prior distribution over rates P(*S=s*) is combined with the likelihood P(*Δg*|*s,w*) to obtain the posterior P(*S=s*|*Δg,w*) at each check. The resulting posterior belief (and derived uncertainty, e.g., entropy) provides a trial-by-trial estimate of inferred progress rate used in subsequent behavioural and neural analyses.

We show that monkeys track and adapt to the latent progress rate. We then identify neural representations of progress rate in both MCC and LPFC, with distinct temporal and functional profiles. MCC expresses a temporally extended long-term goal progress rate signal linked to the long timescale neurons present in the region, but also maintained encoding of feedback information. LPFC expresses progress rate only when inferences on that property were required. Finally, we demonstrate an emergent internal representation of progress rate in hidden units of an RNN trained to predict gauge size. Lesioning progress rate units in the network impairs progress predictions, suggesting progress rate is a critical internal emergent construct in our task.

Together, our results illustrate how MCC–LPFC circuits support progress rate inference and monitoring, providing a neural framework for how prefrontal systems track long-horizon goal progress to guide incremental goal pursuit.

## Results

### Inference of the rate of progress

In this task, the progress rate (the number of correct work trials required per unit gauge increase) is not explicitly cued. Animals must therefore infer the latent rule linking correct discrimination trials/work to bonus progress. Both monkeys rarely checked immediately after incorrect work trials or other check trials (Fig. 1b), indicating that they understood that positive feedback during work drives gauge advancement.

A key question is how animals infer this rate from partial information. The animal observes the gauge only when it checks. A global strategy, such as dividing cumulative correct work trials by cumulative gauge increase, would eventually be accurate but becomes cognitively demanding when blocks are long (up to 36 trials). We therefore hypothesized that monkeys rely on a local, between-check estimate based on consecutive checks: dividing the number of correct work trials, *w*, by the observed gauge increment, *Δg*.

We formalized this local strategy as Bayesian inference over a discrete latent progress rate *s*(Fig. 1c,d,e, see Methods). On each check *n*, the animal observes the gauge increment Δ*g*_*n*_ since the previous check, and (implicitly) has access to work length *w*_*n*_, the number of correct work trials between checks. Because the gauge increases in discrete steps and checks can occur at different positions relative to the unobserved within-step counter, the same observation (*w, Δg*) can be compatible with multiple latent rates *s*. We therefore compute the likelihood *P(Δg*|*w,s)* by enumerating all check positions consistent with discrete gauge-growth under rate *s*, assuming checks are uniformly distributed across positions (Eq. 1). This yields an analytically tractable mapping from (*w, Δg*) to evidence for each candidate rate.

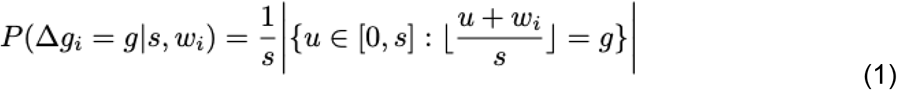

The posterior over progress rates is then updated at each check via Bayes’ rule:

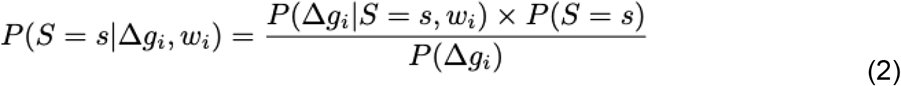

with the prior *P(S = s)* given by the posterior from the previous check.

We summarize the animal’s inferred progress rate at check *n* using the posterior and quantify uncertainty using Shannon entropy. As the number of checks increases within a block, the posterior progressively concentrates around the true rate (Extended Data Fig. 2a), and uncertainty decreases (Extended Data Fig. 2b). Importantly, estimation accuracy is not strictly monotonic: short work lengths between checks are less informative and therefore produce broader posteriors (Extended Fig. 1, 2c), transiently increasing uncertainty even late in a block.

Because the true progress rate is latent and this model provides the optimal estimate under partial observability, we use the model-derived progress-rate belief for subsequent behavioural and neural analyses (except in the RNN simulations, where progress rate is inferred implicitly by the network).

### Behaviour

The Bayesian model provides a normative estimate of the latent progress rate from partial observations. To test whether progress rate actually influences the behaviour of the animals, we examined how it modulates two components of task performance: checking, which trades off information against time cost, and work, which reflects motivation in the perceptual decision task.

#### Progress rate modulates checking behaviour

Checking serves two purposes: it reveals current progress (gauge size) and, when the gauge is full, delivers the bonus. However, frequent checking is time consuming, since closely spaced checks yield less new information (Extended Data Fig. 1c), and displace work trials that could have produced immediate juice reward. Under slower progress rates, it is therefore strategically advantageous to space checks further apart.

To evaluate whether monkeys adaptively adjust check spacing, we plotted the average distance to the next check (number of intervening work trials) as a function of current gauge size, separately for different progress rate conditions (colour-coded; Fig. 2a). Two patterns emerged. First, regardless of rate, as the gauge approached its maximum, both animals sharply reduced the distance to the next check, consistent with a drive to secure the bonus as soon as it became available. Second, check spacing depended on progress rate, but in an animal-specific manner. Monkey A showed the expected pattern: checks were spaced further apart under slower progress. Monkey H showed an opposite trend. This inversion plausibly reflects a distinct strategy in the fastest-rate condition, in which the animal worked long enough to get the bonus without checking intermittently then timed one check to capture it directly.

**Figure 2.**
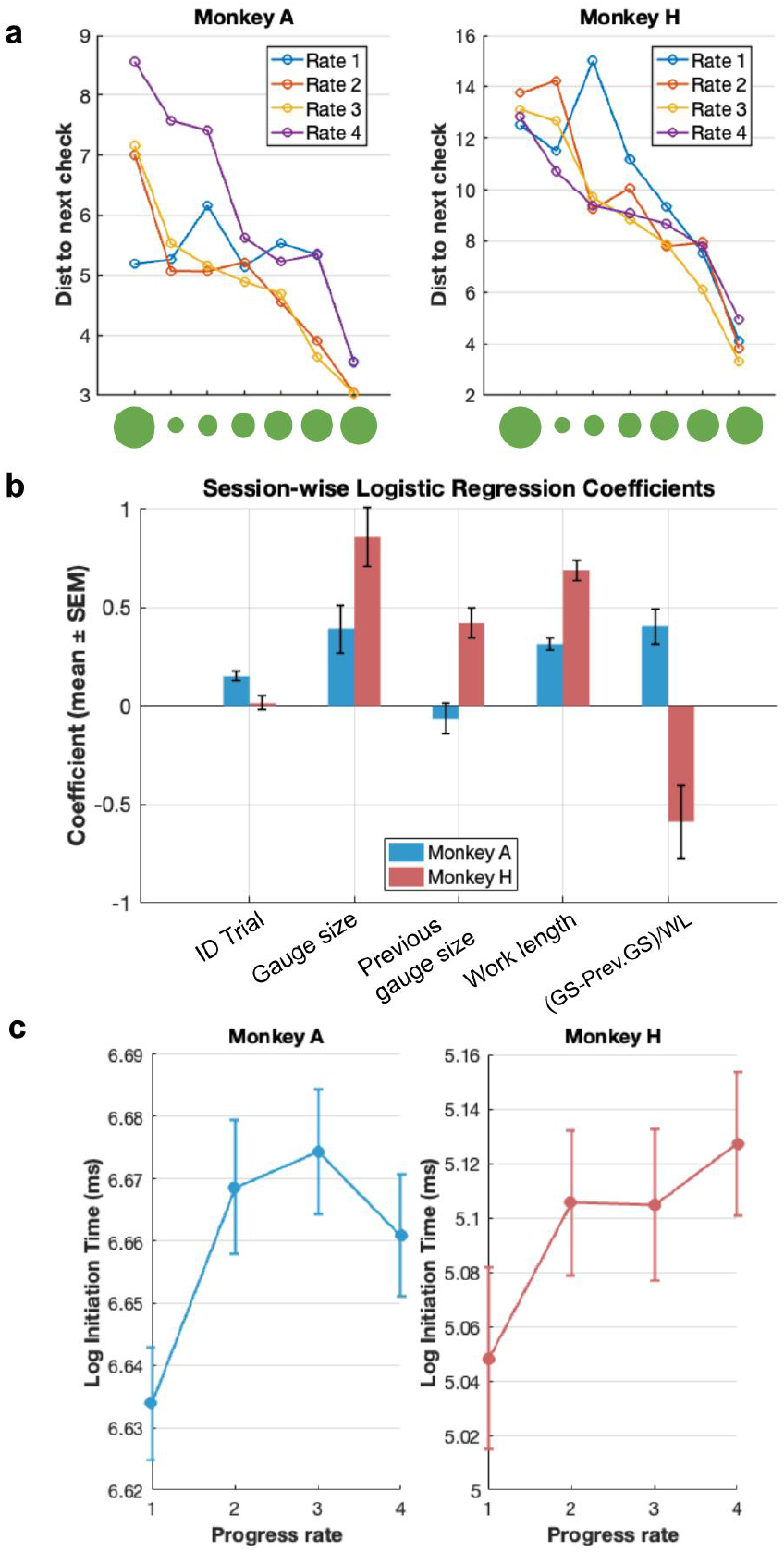
Behavioural evidence of progress rate sensitivity. **(a)** *Check spacing depends on gauge size and progress rate*. Mean distance to the next check (number of intervening work trials) plotted against current gauge size, separately for each progress-rate condition, shown for Monkey A (left) and Monkey H (right). Across rates, check spacing decreases as the gauge approaches full, consistent with increasing checks to capture the bonus. *Progress-rate contributes significantly to predictions of checking*. Session-wise logistic regression coefficients (mean ± SEM across sessions) predicting the probability of choosing check on each trial from trial index (time-on-task), current gauge size (GS), gauge size at the previous check (Prev. GS), work length defined as the number of correct work trials since the previous check (WL), and a local progress-rate evidence term (GS−Prev. GS)/WL. Coefficients are shown separately for each monkey (colours) with z-scored regressors, allowing comparison of effect magnitudes. **(c)** *Progress modulates initiation time*. Mean log initiation time in work trials as a function of progress rate for Monkey A (left) and Monkey H (right), indicating systematic slowing under slower progress (error bars indicate variability across sessions/blocks).

Because check spacing covaries with proximity to the bonus, we next asked whether progress rate predicts checking after controlling for gauge size and work length. We fit a logistic regression predicting the probability of checking on each trial from: trial number (capturing time-on-task), gauge size at current and previous check, work length, and a local estimate of progress-rate. All regressors were z-scored to enable coefficient comparison.

As shown in Fig. 2b, checking in both monkeys was dominated by current gauge size and work length, consistent with a strategy primarily focused on bonus timing and proximity. Trial number contributed modestly, and only for Monkey A, consistent with a weak time-on-task effect. Crucially, the local estimate also significantly predicted checking in both animals, indicating that animals used progress rate information beyond the current gauge size. The direction of this effect differed between animals, mirroring the check-spacing analysis: Monkey A showed the direction expected from an optimal spacing strategy, whereas Monkey H showed the opposite sign, consistent with its alternative “work-till-the-end” strategy in the fast-rate regime.

#### Progress rate shapes motivation to work

If animals track progress rate, it should also influence how fast they initiate work. We therefore examined initiation times on work trials (time between lever onset and choosing to work) as a function of progress rate(Fig. 2c). For both monkeys, initiation times slowed under slower progress, consistent with reduced motivation when progress is slow. A linear regression confirmed significant effects of progress rate on initiation time (linear mixed-effects model: LogIT ∼ ProgressRate + (1|Monkey), *β* = 0.0130 ± 0.0030, *p* = 1.67e-5).

Together, three converging analyses, (i) distance to the next check, (ii) logistic regression controlling for gauge size, time-on-task, and recent history, and (iii) initiation-time modulation, show that behaviour in both animals is sensitive to progress rate. Although checking is strongly driven by bonus acquisition, progress rate nonetheless measurably shapes both information-seeking and motivation, consistent with animals maintaining and using an internal estimate of the latent action-to-progress mapping.

### Learning progress rate at check trials

Having established behavioural sensitivity to progress rate, we next asked whether behavioural and neural activities (Fig. 3a, see Methods for neural recording details) reflect the inference process. Because beliefs about progress rate are updated primarily at check trials when the gauge is revealed, we focused analyses on the choice time window surrounding check decisions.

**Figure 3.**
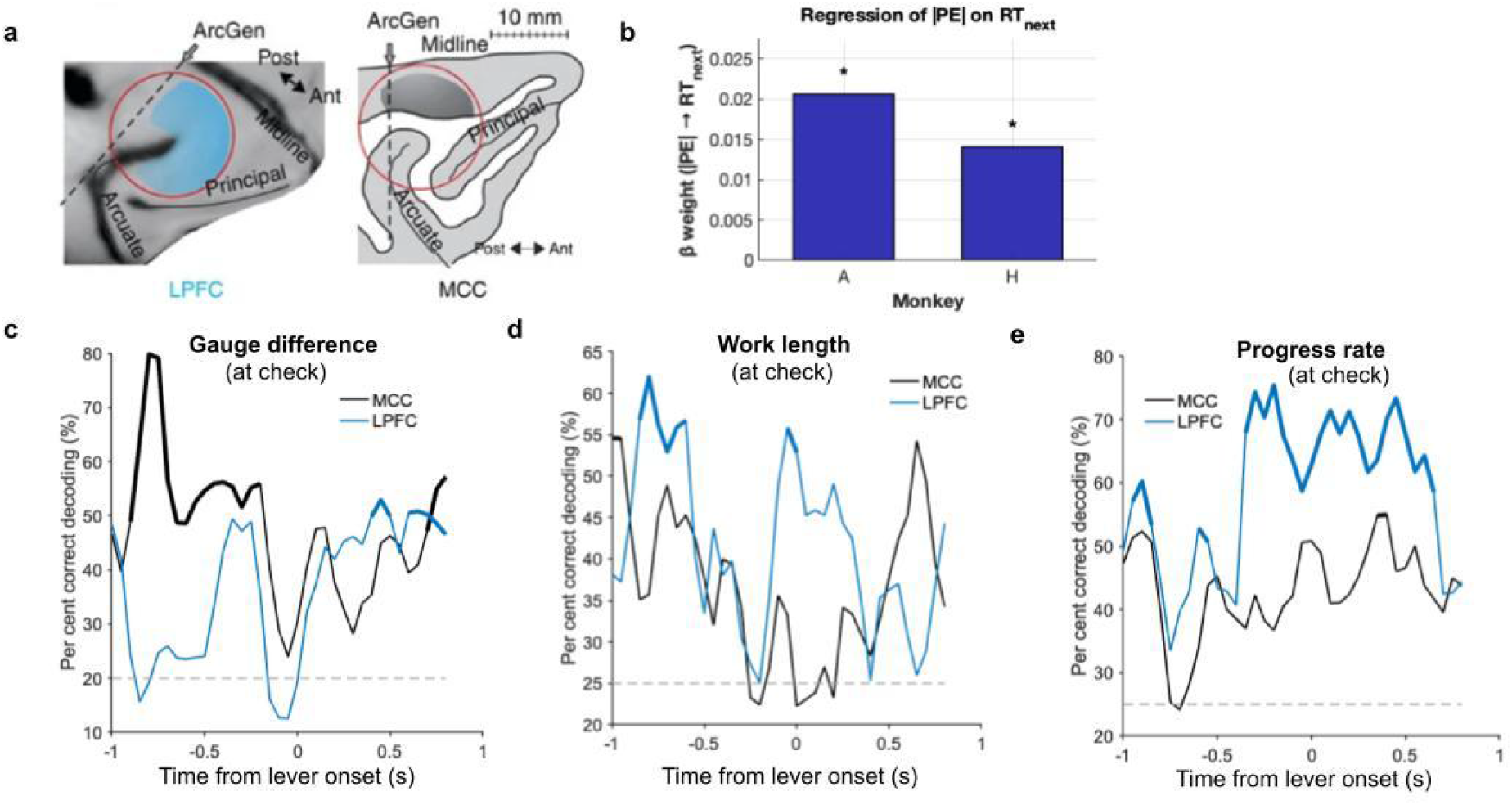
Learning progress rate at checks: neural correlates of Bayesian evidence. **(a)** *Recording sites*^7^. Example recording locations in lateral prefrontal cortex (LPFC; left) and midcingulate cortex (MCC; right) from the two monkeys. **(f)** *Behavioural impact of progress prediction error at the next work trial*. Regression weight relating progress prediction error (PE; derived from the Bayesian model) to subsequent behaviour (next-trial reaction time), shown separately for each monkey. Significant weights (asterisks; t-test) indicate that surprise at checks slows down the next work trial, suggesting an update of the progress-rate estimation. **(c)** *Neural representation of gauge evidence*. Time-resolved decoding accuracy for the gauge increment *Δg* between consecutive checks (a key Bayesian evidence term), using a sliding-window linear SVM classifier on pseudo-population firing rates. MCC (black) and LPFC (blue) are shown; green dashed line indicates chance. Thicker lines indicate significance by permutation test. **(d)** *Neural representation of work-length evidence*. Time-resolved decoding accuracy for work length *w* (number of correct work trials since the previous check), plotted as in (b). **(e)** *Neural representation of inferred progress rate*. Time-resolved decoding accuracy for progress rate during the same choice window on **check trials**, demonstrating robust rate information in LPFC.

#### Prediction errors at checks slow subsequent work

At each check, the animal can compare the gauge size it would expect based on its current estimate of the progress rate (and the known work length) to the observed gauge size. The difference between the expected and observed gauge size provides a prediction error that can be used to update the rate estimate. We therefore quantified this check-related prediction error and asked whether it influenced subsequent behaviour. A regression of the absolute prediction error against reaction time on the next work trial revealed that larger prediction errors were followed by longer reaction times (Fig. 3b, t-test, monkey A: *β* = 0.0207, *p* = 0.0012; monkey H: *β* = 0.0180, *p* = 2.4e-5), consistent with the idea that surprising belief updates transiently slow task execution as animals incorporate new information into an internal model of progress.

#### Pre-choice representations of the elements for inference

We first decoded the key variables required by the Bayesian update: the work length *w* (number of correct work trials since the previous check, discretized into 4 equal-percentile bins by pooling trials within each monkey), gauge increment *Δg* (the change in gauge level across checks, ranging from 0 to 6) using pseudo-population activities during only check trials (see Methods for details). All variables were significantly decodable above chance before the check-versus-work choice was made (Fig. 3c,d). This indicates that, when animals choose to check, monkeys maintain internal information sufficient to support progress-rate inference. Consistent with prior work, the two regions showed complementary encoding dynamics: MCC led in encoding gauge-related information (goal proximity), whereas LPFC led in encoding work-length information (parameters of ongoing task execution)^7^. This dissociation suggests a division of labor in which LPFC preferentially maintains the recent action history needed to interpret a check outcome, while MCC preferentially tracks progress toward the bonus.

Together, these results provide convergent evidence that progress rate is inferred at checks: pre-choice activity contains the information required for inference, transforming into activity reflecting more overall progress rate.

### Neural representation of progress rate

We have shown that progress rate is decodable during check trials, when animals explicitly reveal the gauge and update predictions. An open question is whether this signal is transient, emerging only during gauge prediction and reveal, or whether it functions as a latent contextual variable that persists across the block and conditions other computations, including feedback processing during work.

To address this, we extended our time-resolved decoding analyses to include both work and check trials, and centered analyses not only on choice epochs but also on the feedback window of the main task. Progress rate was decodable well above chance in both MCC and LPFC across trial types and around feedback (Fig. 4a,b, see Methods for details). Importantly, decoding remained robust throughout extended portions of these epochs rather than appearing as a brief burst locked to a single event, consistent with a tonic, context-dependent representation of inferred progress rate. Across epochs, MCC decoding exceeded LPFC decoding, suggesting that progress-rate information is represented more precisely and more consistently in MCC.

**Figure 4.**
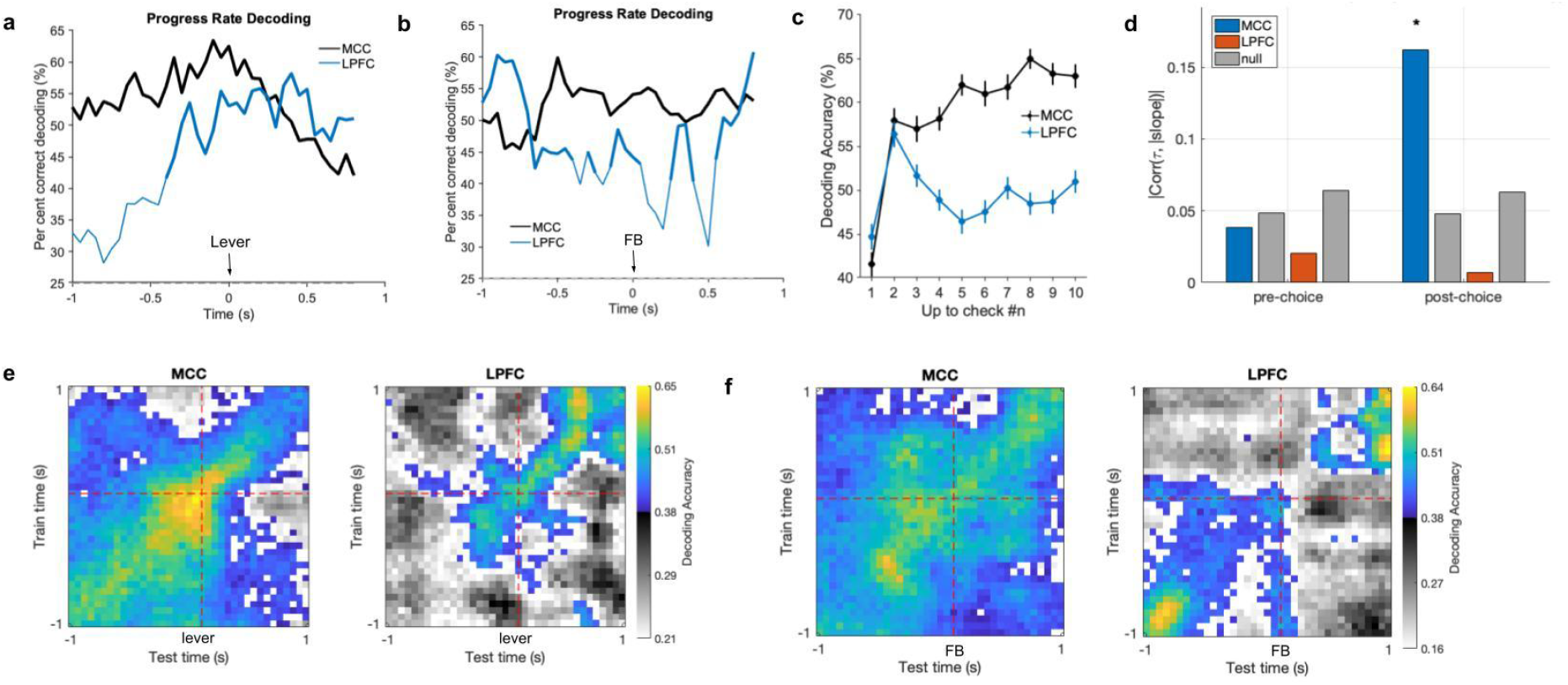
Progress rate is represented as a sustained contextual signal and relates to intrinsic neuronal timescales. **(a)** Time-resolved decoding accuracy for progress rate during the choice window, shown separately for MCC (black) and LPFC (blue) with significant decoding performance marked by thicker lines (permutation test). The green dashed line indicates chance. **(b)** Time-resolved decoding accuracy for progress rate during the feedback window, plotted as in (a). **(c)** Across-check refinement of rate information. Progress-rate decoding accuracy evaluated at work/check decision as a function of check number within a block (“up to check #n”), showing rapid improvement after early informative checks followed by more gradual refinement only in MCC (mean ± SEM). **(d)** Correlation between ***τ*** and progress-rate encoding magnitude computed in fixed pre- and post-feedback windows, shown for MCC and LPFC. Asterisk indicates a significant post-feedback association in MCC (two-sided t-test). **(e)** Cross-temporal decoding matrices for MCC (left) and LPFC (right) around choice, obtained by training the decoder at one time point (y-axis) and testing at all other time points (x-axis). MCC shows broad generalization across time (stable format), whereas LPFC generalization is more time-specific. **(f)** Temporal stability of rate coding around feedback (cross-temporal generalization), shown as in (c). Again, MCC exhibits stronger off-diagonal generalization than LPFC, consistent with a more time-invariant representational format.

Importantly, decoding accuracy also increased across successive checks and closely mirrored the sharpening of the Bayesian posterior, particularly in MCC (Fig. 4c). Notably, decoding accuracy in LPFC did not increase with more checks, indicating that the improvement in MCC is not a trivial consequence of having more trials available for decoding. This pattern supports the interpretation that monkeys refine a latent progress-rate belief through accumulation of evidence over checks.

We next asked whether the underlying neural code is temporally stable. Using cross-temporal generalization (training at one time point and testing at others), we found a marked regional dissociation. In MCC, classifiers generalized strongly across time within both choice and feedback windows, indicating a stable population pattern for progress-rate encoding that persists across task moments. In contrast, LPFC exhibited more event-linked coding: cross-temporal generalization declined rapidly outside the training time point, consistent with a weaker and dynamic representation whose format changes across epochs and is tied to specific task events.

This MCC–LPFC dissociation suggests that inferred progress rate serves as a block-level contextual state rather than a transient “check-only” computation. In MCC, strong cross-temporal generalization is consistent with a stable population code that can continuously parameterize how outcomes should be interpreted and how vigorously to act, effectively providing a running contextual scaffold for sequential decisions and feedback processing. By contrast, LPFC’s weaker temporal generalization is consistent with a dynamic, event-linked format that is well suited for selectively updating beliefs and policies when informative evidence arrives (for example, around choice preparation or immediately following feedback), rather than maintaining a fixed code throughout the trial. Such phasic or dynamic population codes are widely observed in prefrontal cortex and are thought to support flexible computation while still enabling accurate readout of task variables^7,18–21^.

### Timescales of neurons carrying progress rate information

The decoding analyses above point to two complementary regimes for representing progress rate: a stable, context-like signal in MCC that generalizes across time and trial events, and a more event-locked, dynamic signal in LPFC. This dissociation raises a mechanistic question: what intrinsic properties of neurons support progress-rate information, given that inferring the block-wise rate requires integrating evidence across trials?

One candidate is intrinsic timescale (time constant, *τ*), estimated from the decay of spike-count autocorrelation. Across cortex, intrinsic timescales are hierarchically organized, with association regions exhibiting longer timescales than sensory regions, consistent with a greater capacity for temporal integration^22^. Computational work suggests that such hierarchical timescales can emerge from differences in recurrent circuit properties^23^. In the frontal cortex specifically, MCC has been reported to exhibit longer timescales than LPFC and to express temporal signatures linked to metastable network dynamics, consistent with a role in integrating outcome history across trials^24^.

To test whether intrinsic timescale relates to progress rate coding in our task, we regressed each neuron’s firing rate against progress rate and quantified progress rate “relevance” as the absolute regression slope. We then correlated this value with each neuron’s intrinsic timescale. Following prior work on intrinsic timescale estimation^22,21,25,26^ to reduce sensitivity to unstable autocorrelation fits, we excluded neurons with *τ* > 900 ms (Extended Data Fig. 3a, see Methods).

During the feedback window, the time-resolved correlation between *τ* and progress rate regression magnitude increased after feedback onset and then returned toward baseline only in MCC (Extended Data Fig. 3b). In fixed windows (500 ms pre-vs. 500 ms post-feedback), the correlation was significant only in MCC after feedback (Fig. 4d; Pearson correlation *r* = 0.162, permutation test: *p* = 0.0285). Together, these results align MCC’s role in monitoring and updating latent progress dynamics from outcomes with a selective contribution of longer-timescale neurons precisely when information must be integrated to update state estimates across trials.

During the choice window, we found no reliable relationship between intrinsic timescale and progress rate regression magnitude in either MCC or LPFC (Extended Data Fig. 3c). However, MCC showed a non-significant transient post–gauge reveal peak (Extended Data Fig. 3d), consistent with the idea that longer-timescale neurons may be preferentially engaged at moments when newly revealed information can update the inferred progress state.

### Feedback geometry across progress rates

A correct work trial does not carry the same implications for future progress under different progress rates: the mapping from outcome (feedback) to gauge advancement changes with the latent rate. Motivated by the timescale results suggesting that progress rate information is recruited around feedback, we asked whether progress rate modulated the neural population geometry of feedback in a specific way to support flexible readout across rate contexts. One possibility is that neural dynamics of opposite feedback shift parallelly when progress rate changes, allowing an extraction of both generalized feedback information and progress rate specific information, which could be used to update progress.

To visualize how feedback valence and progress rate are mixed, we projected the mean population responses onto the first three principal components (PCs) (Fig. 5a). In both MCC and LPFC, feedback-evoked activity patterns were separated across progress rates, indicating that rate modulates outcome representation. However, this modulation differed between areas. In MCC, the direction separating positive from negative feedback appeared largely preserved across progress rates. In LPFC, by contrast, the positive-versus-negative separation rotated more strongly across progress rates, suggesting a greater mixing of feedback valence with latent progress rate.

**Figure 5.**
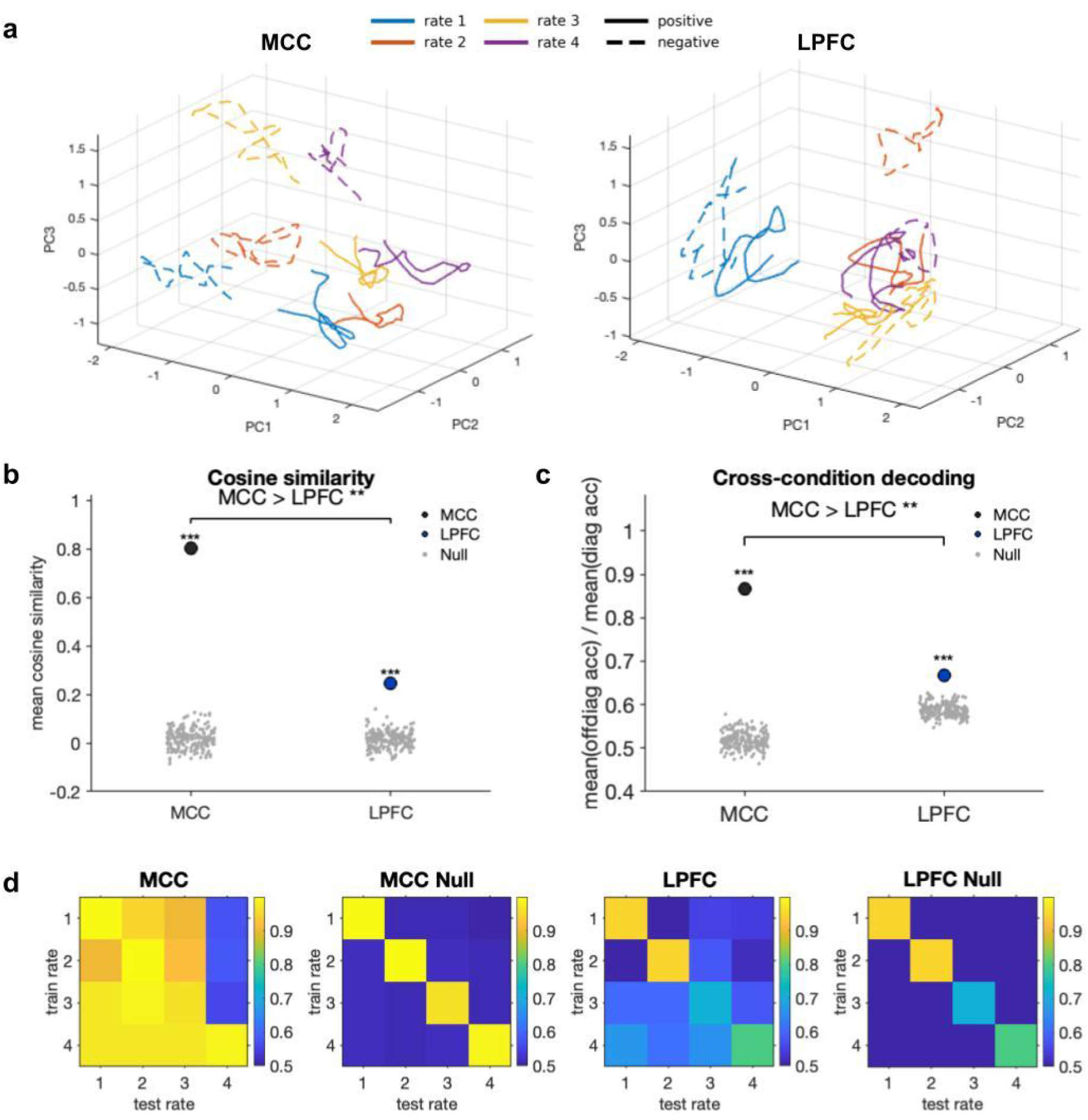
Progress rate modulates feedback responses while preserving a stable valence axis in MCC. **(a)** *Low-dimensional visualization of feedback axes across progress rates*. Trajectories of mean population activity projected onto the first three principal components, shown separately for MCC (left) and LPFC (right), with colours indicating progress-rate condition. Feedback responses are separated by progress rate context in both areas, but MCC maintains a more consistent axis between positive and negative outcomes across contexts. **(b)** *Alignment of feedback axes across progress rate contexts*. Mean cosine similarity of the positive-negative feedback vectors computed for all pairs of progress-rate conditions, compared against the null distribution. MCC exhibits higher axis alignment (greater parallelism) than LPFC, indicating a more context-invariant valence axis. **(c)** *Feedback generalization exceeds null and is stronger in MCC*. Summary metric of cross-condition generalization (normalized relative to within-condition decoding; as defined in Methods) compared against the null distribution for each region. Coloured dots indicate observed values; gray points show the null performance; bracket denotes a significant MCC > LPFC difference. **(d)** *Cross-condition feedback decoding and null control*. Confusion matrices showing cross-condition decoding accuracy for feedback valence (positive vs negative) when training the classifier in one progress-rate condition and testing in another, shown for MCC and LPFC (top). Corresponding null matrices (bottom) are obtained after shuffling neuron identity, yielding near-chance generalization.

We quantified this alignment using the cosine similarity between the positive-minus-negative feedback vectors for all pairs of rate conditions. Both regions exhibited cosine similarities significantly above the null distribution, indicating shared feedback structure (Fig. 5b; *p* = 5e-4). Importantly, MCC showed significantly higher parallelism than LPFC (Fig. 5b; *p* = 0.003), consistent with a generalized feedback axis relatively invariant to progress rate context. This pattern suggests that progress rate modulates feedback activity without substantially rotating the core positive-versus-negative feedback dimension in MCC, yielding an outcome representation that is more disentangled from latent rate context^27,28^.

We also quantified feedback-axis alignment using cross-condition decoding of feedback. Classifiers were trained to discriminate positive versus negative feedback under one progress-rate condition and then tested either within the same condition or on a different condition. In both MCC and LPFC, within-condition decoding reached high accuracy (typically >90%), whereas cross-condition performance decreased, indicating rate-dependent modulation of feedback responses (Fig. 5c,d). Cross-condition decoding nevertheless remained significantly above chance (*p* = 5e-4) in both regions when compared to a null distribution generated by permuting unit identities across conditions, which reduced cross-condition performance to near chance (50%) while preserving overall firing-rate structure (Fig. 5c,d). Remarkably, MCC generalized significantly better than LPFC across rate conditions (Fig. 5c; *p* = 0.003), suggesting that the feedback representation is more conserved across contexts in MCC.

Together, these results reveal complementary feedback codes. LPFC expresses feedback in a more context-dependent format, whereas MCC preserves a more context-invariant feedback direction, providing a stable outcome-evaluation signal that can be read out consistently across blocks even as progress-rate context changes.

### Progress rate representations emerge naturally in RNN hidden units and are critical to predict progress

The analyses above show that monkeys infer a latent progress rate and that this contextual variable shapes neural activity, including how feedback is represented across conditions. What remains unclear is what computational pressure is sufficient for a progress rate signal to emerge. A natural candidate is the need to predict covert progress without constant checking. To test this, we trained a recurrent neural network (RNN) to predict gauge size from the same partial action–outcome history available to the animals (Fig. 6a). This framework also supports causal perturbations of units with rate-related representations.

**Figure 6.**
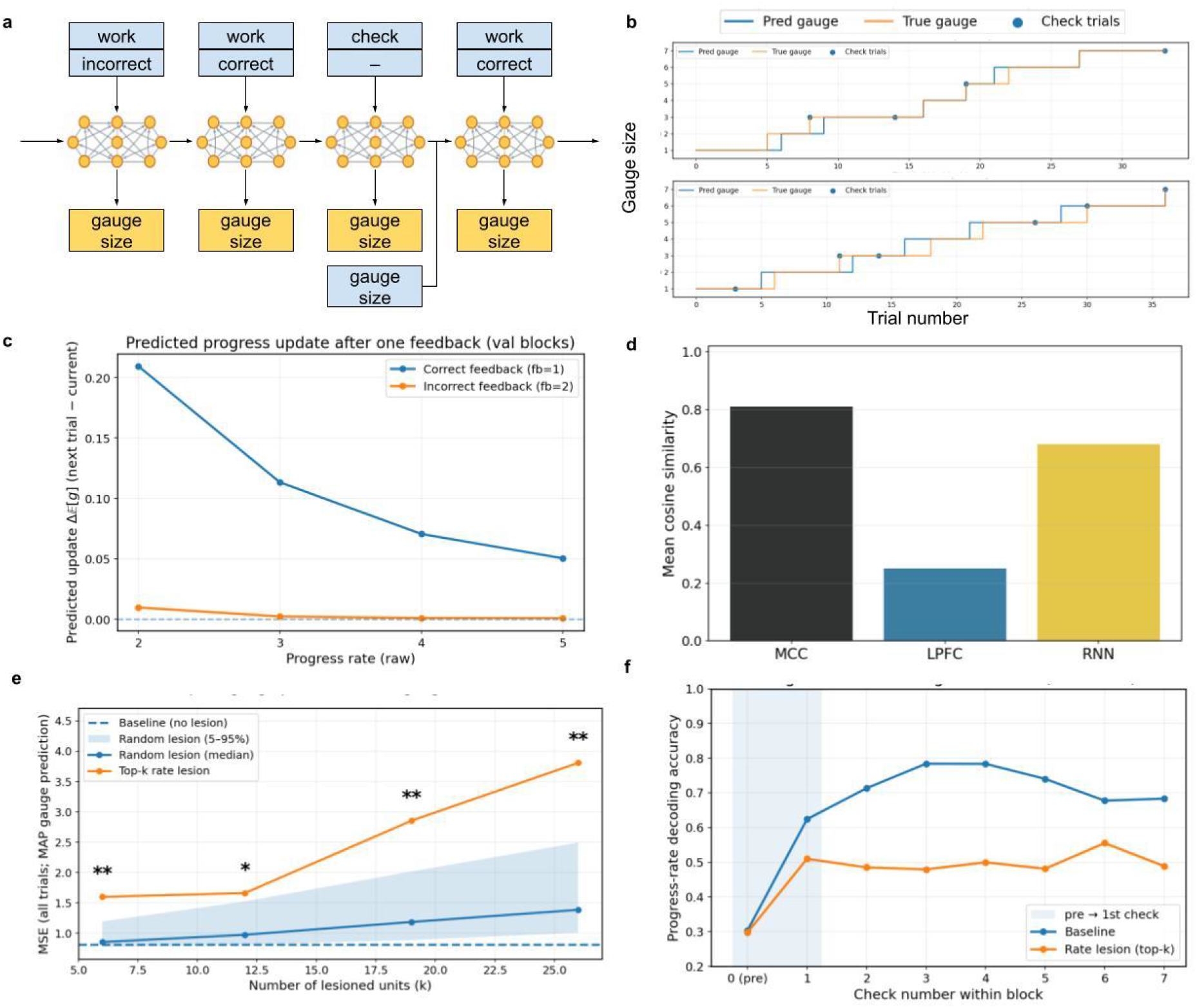
A recurrent neural network trained to predict progress develops progress-rate representations that support accurate gauge tracking. **(a)** *RNN schema and training regime*. Schematic of the recurrent neural network receiving the same partial trial history available to the animals: choice (work vs check) and feedback (correct/incorrect) on work trials, with gauge value revealed only on check trials. The network outputs a running prediction of gauge size each trial; loss is computed only on check trials, where the predicted gauge is compared to the observed gauge. On check trials, the true gauge is provided after the prediction and used only to update the recurrent state for subsequent trials, preventing information leakage. **(b)** *Accurate gauge prediction from intermittent observations*. Example validation blocks showing the true gauge trajectory (orange) and the RNN’s predicted gauge (blue) across trials, with check trials marked. The network tracks gauge dynamics across trials despite sparse supervision. **(c)** *Learned progress update rule*. Predicted gauge update following feedback as a function of progress rate condition: correct feedback produces a positive increment whose magnitude scales with rate, whereas incorrect feedback produces little or no increment, indicating that the network learned the task contingency linking correct work to progress. **(d)** *Abstract feedback geometry in the RNN*. Mean cosine similarity between positive-negative feedback axes across all pairs of progress-rate conditions, shown for MCC, LPFC, and the RNN model. High axis alignment in the RNN indicates a relatively rate-invariant feedback encoding, resembling the context-general feedback geometry observed in MCC. **(e)** *Targeted lesion of “rate units” impairs progress prediction*. Mean squared error of gauge prediction as a function of the number of lesioned units *k*, comparing baseline (no lesion), random lesions, and targeted lesions of the top-*k* “rate units” (identified by a linear regression of progress rate from hidden activity at checks). Asterisks denote significant differences between targeted and random lesions. **(f)** *Progress rate representations emerge and sharpen across successive checks, and targeted lesions disrupt this refinement*. Progress rate decoding accuracy from hidden activity evaluated at checks as a function of check number within a block, shown for the intact network and the targeted-lesion network (k=26). Shaded region marks the abrupt post–first-check increase in decoding accuracy, a putative signature of inference.

On each trial, the network received the animal’s choice (work vs. check), trial feedback when applicable, and gauge value only on check trials. The network produced a running prediction of gauge size, but supervision was sparse: the loss was computed only on check trials as the mean squared error between predicted and observed gauge size (Fig. 6a). To encourage stable trajectories, we included an auxiliary smoothness loss on consecutive hidden states, and to enforce the task contingency we penalized prediction updates following incorrect work. Critically, on check trials the observed gauge size was revealed only after the network generated its prediction and was used only to update the recurrent state for subsequent trials, preventing trivial copying. Under this regime, the RNN accurately tracked gauge dynamics across blocks, demonstrating that progress can be inferred from intermittent checks and trial-by-trial feedback (Fig. 6b).

Because the model produces trial-wise gauge predictions, we could also examine feedback-driven updates under different rates. The predicted gauge increment following correct feedback scaled with rate, whereas incorrect feedback produced almost no increment (Fig. 6c), demonstrating that the network learned that progress depends on correct work trials and grows linearly at a block-specific rate. Besides, feedback decoding axes of the RNN exhibited high cosine similarity across rate pairs (Fig. 6d), indicating an abstract feedback representation disentangled from rate estimate, resembling the geometry observed in MCC.

If accurate progress prediction requires inferring the latent progress rate, the network’s internal state should contain a progress rate signal that sharpens with evidence. To test this, we extracted the hidden state at each check and decoded the block’s progress rate. Progress rate was reliably decodable, with accuracy rising sharply after the first informative check and improving more gradually thereafter (Fig. 6f), mirroring the progressive sharpening of the Bayesian estimate and the across-checks neural decoding results (Fig. 3e).

We next asked whether rate-related activity is functionally necessary for accurate progress prediction. We identified “rate units” by fitting a linear model to predict progress rate from hidden activity at checks and ranking units by the absolute regression weight. Lesioning increasing fractions of the top-ranked units (∼5%, 10%, 15%, 20% of 128) by clamping their activity to zero while leaving all network weights unchanged increased gauge-prediction error relative to random-lesion controls (Fig. 6e; p(k=6)=0.0065, p(k=12)=0.0345, p(k=19)=0.0090, p(k=26)=0.0035). Despite this impairment, progress rate decoding from the lesioned network remained above chance (Fig. 6f), indicating that rate information persisted in the remaining dynamics. Notably, however, the lesioned network showed reduced refinement of the rate representation across successive checks, showing the importance of the lesioned units that maintain a stable, context-like rate signal across trials.

Together, these simulations provide a computational complement to the neural data: when progress must be inferred from intermittent checks, a progress rate signal emerges naturally in recurrent state, refines with experience, and supports efficient progress tracking. This offers a mechanistic account of how rate inference and abstract feedback coding can arise from the core behavioural problem: securing the bonus quickly while avoiding excessive, uninformative checking.

## Discussion

Long-term goal pursuit requires estimating how fast actions translate into future progress, particularly when progress rate changes across goals or progress observations are only sparse. Here we show that macaques infer a latent progress rate from sparse observations, and that this inferred variable impacts neural activity across MCC and LPFC, but in different ways: MCC carried a comparatively stable signal that generalized across time and trials, whereas LPFC activity was linked to progress rate transiently, in particular during initial progress rate inference. The MCC signal was consistent with a role in rate inference, progress tracking and feedback processing once the actual rate was known. Importantly, progress rate also changed how outcomes were processed, modulating basic feedback responses. While a positive–negative feedback axis was particularly conserved in MCC, LPFC mostly contained progress rate specific feedback signals. Finally, an RNN trained under the same partial observability constraints developed a progress rate representation that refined with evidence and was functionally important for accurate progress prediction, demonstrating that rate-like signals and stable feedback geometry can emerge naturally from the demands of tracking covert progress from intermittent checks.

A central conceptual result concerns the geometry of feedback modulation by progress rate. In this task, identical outcomes carry different implications for future progress depending on the inferred progress rate. Nevertheless, cross-condition decoding and axis-alignment analyses revealed a substantial component of feedback coding that generalizes across rate contexts, especially in MCC. This links progress monitoring to recent work on abstraction and representational geometry in high-dimensional neural populations^28–30^. In the framework developed by Fusi and colleagues, abstract variables are characterized not merely by decodability within a condition, but by a geometry that supports linear generalization across contexts, often quantified via alignment (“parallelism”) of coding directions across conditions. In our data, the positive-negative feedback axis remained comparatively aligned across progress-rate contexts in MCC, supporting cross-condition generalization, whereas LPFC exhibited greater axis rotation and weaker generalization. This suggests that progress rate context modulates feedback-evoked activity in a subspace orthogonal to feedback encoding axis in MCC, offering a population-level mechanism by which outcome valence can remain consistently readable even as the meaning of outcomes for future advancement depends on a changing latent context.

The distinct neural patterns in MCC and LPFC were further consistent with general functional dissociations. A stable representation of progress rate and integration of feedback signals could support monitoring progress and determine behaviours such as vigor across extended sequences of trials, as well as contextualize behaviour by progress rate when explicit progress observations are sparse. In contrast, a more transient signal in LPFC may be advantageous for selectively updating beliefs or policies in noisy environments when new information arrives (e.g., around checks) and when contexts need to be inferred or change across trial epochs. Importantly, these interpretations are grounded in multiple observations: Firstly, MCC showed stronger cross-temporal generalization, indicating that progress-rate information was available in a relatively time-invariant format across task moments, whereas LPFC’s reduced temporal generalization indicates a more event-linked format that may prioritize flexible, time-specific computation. Secondly, LPFC signals linked to inference relevant information (work length and gauge change) mostly emerged at checks, suggesting more selective recruitment of this circuit during novel observations. Lastly, when actual progress towards the goal was made, albeit at variable rates, LPFC also had weaker progress rate signals. However, MCC appeared to have another role in further signalling the general effect of success and failure of work, suggesting MCC’s importance in not just monitoring and updating progress for its impact on behaviour, choice and motivation but also abstracting away from any current rate of gains to signal the general success of failure of an action.

Our intrinsic-timescale analyses provide a mechanistic constraint on how MCC might achieve its multiple hypothesized functions. Updating an inferred progress rate requires integrating evidence across trials, and intrinsic timescales have been proposed as a signature of temporal integration^24^. We found that the relationship between intrinsic timescale and progress-rate encoding was temporally specific, emerging most clearly following feedback in MCC. This timing is notable: feedback is precisely when new evidence should update latent progress dynamics. The association between longer timescales and stronger post-feedback progress-rate encoding suggests that slow, integrative neural subpopulations contribute selectively when outcome information must be accumulated to update a context variable that persists across trials. While correlational, this result motivates a testable hypothesis: perturbations that selectively disrupt long-timescale subpopulations should disproportionately impair the maintenance or refinement of progress-rate representations, particularly in MCC.

The RNN simulations complement our physiological results by isolating a minimal computational pressure sufficient to generate progress rate representations. The network was trained to predict gauge size under sparse supervision (only at checks), and thus faced the same core challenge as the animals: progress must be tracked covertly from intermittent observations at checks and trial-by-trial feedback. Under this constraint, progress rate became decodable from hidden state and sharpened with successive checks, mirroring both the Bayesian estimate and the neural decoding trajectory. Targeted lesions of “rate units” increased prediction error significantly more than random lesions, indicating that rate-related internal structure was functionally important for accurate progress tracking. Interestingly, progress rate decoding remained above chance after lesioning, but refinement across checks was diminished, consistent with different subcomponents of the recurrent dynamics contributing to prediction from initial observation versus further refinement using stable maintenance across trials. Together, these simulations support the view that progress-rate representations arise naturally when a system must predict latent progress from incomplete information, and they can confer robustness and efficiency with repeated observations or long-horizon behaviour.

Overall, our findings identify latent progress rate as a key variable that shapes behaviour and neural representations during long-term goal pursuit. This is differentially supported across the prefrontal cortex: LPFC carries transient, event-locked signals consistent with rate inference at checks, whereas MCC maintains a stable, cross-temporally generalizable rate representation that supports continuous monitoring and feedback-driven updating. Future work can build on the causal leverage of our RNN framework and on prior evidence for hierarchical cortical dynamics^24^ by moving beyond discrete-time implementations to models that explicitly represent neural timescales and inhibitory control. In particular, continuous-time RNNs and more biophysically grounded circuit models will allow intrinsic timescale estimates that are directly comparable to empirical measures, enabling a stringent test of whether longer time constants preferentially support stable context maintenance and refinement of progress-rate beliefs. These models will also examine how inhibition and timescale interact to produce area-specific computations, including MCC-like stability versus LPFC-like event specificity, and whether targeted perturbations of long-timescale or inhibitory subpopulations selectively disrupt the construction and maintenance of progress rate information.

## Methods

### Dataset

Two male rhesus monkeys (*Macaca mulatta*), referred to as A and H, were trained to perform a *check vs. work* task while single-unit activity was recorded from the midcingulate cortex (MCC) and lateral prefrontal cortex (LPFC). At the beginning of each trial, the monkeys could choose to either engage in a visual categorization task (*work*) or inspect their progress toward a larger reward (*check*).

In *work* trials, the monkeys were required to identify and select the target stimulus that matched the orientation of a previously shown cue. Correct responses were immediately rewarded with juice (positive feedback), incorrect responses were followed by an absence of reward (negative feedback). In *check* trials, a visual gauge was displayed on the screen, indicating progress toward a large bonus reward equivalent to seven times the reward of a correct work trial. This gauge increased incrementally with each correct work trial and is shown only when the monkey chose to check it. Once the gauge reached its maximum size (7 units) and the monkey opted to check, the accumulated bonus was delivered, marking the end of a block.

Importantly, the rate at which the gauge increased varied across blocks: the number of correct work trials required to earn the bonus was 14, 21, 29, and 36 for progress rates 1, 2, 3, and 4 respectively. This manipulation prevented the monkeys from relying solely on internal trial counting. To accurately estimate their progress and obtain the bonus, they were required to intermittently check the gauge rather than waiting passively for a presumed completion point.

For additional details on neural recordings and task design, see Stoll et al.^7^.

### Bayesian inference model

The optimal estimation of the progress rate given the observation of the change of gauge size and the work length can be modeled by a Bayesian model. With the same notation as in the Results section, we can compute the posterior probability given one pair of observation (Δ*g*_*i*_, *w*_*i*_) using

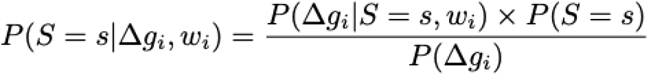

The initial prior is a uniform distribution. Then for check *i*, the prior is the posterior of check *i-1*. Thus given a sequence of observations (Δ*g*_*i*_, *w*_*i*_)_*i*∈{1,… *n*}_, the probability of a certain progress rate is

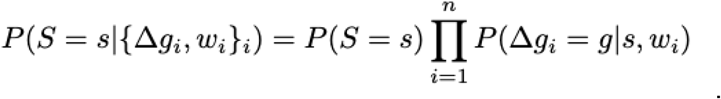

As an intuition-building example of one update step, consider *w=7* correct work trials between checks and an observed gauge increment *Δg=2*. Under rate *s=2* (one gauge step per three correct trials), the first check could occur at three possible phases of the latent growth cycle. Two of these phases yield an observed increment of two steps after *w=7* additional correct trials, thus *P(Δg=2*∣*w=7,s=2)=2/3*. Repeating this calculation for other candidate rates produces a likelihood profile over *s* (Fig. 1d), which is combined with the prior to yield an updated posterior belief (Fig. 1e, Extended Data Fig. 1).

The maximum a posteriori (MAP) estimation of the progress rate is

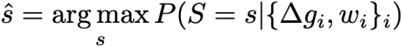

To evaluate the evolution of accuracy of the estimation over checks, the estimation using trials up to *n* checks is computed and compared with the ground truth for each block and the average over blocks is plotted.

The prediction error is defined as the discrepancy between the predicted gauge size and the observed gauge size.

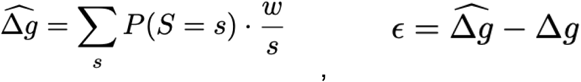

### Neural decoding analyses

All neural analyses pooled single-unit recordings from both monkeys, separately for each brain area (MCC: *n* = 294 units total; 190 from monkey A and 104 from monkey H; LPFC: *n* = 267 units total; 168 from monkey A and 99 from monkey H). For each decoding analysis, we included only neurons that contributed at least one trial to every class of the variable being decoded (i.e., neurons missing one or more classes for a given classification problem were excluded from that problem). Firing rates are computed on non-overlapping bins of 200 ms without smoothing or preprocessing.

Pseudo-population samples were generated as follows. For each decoding target (progress/gauge size, progress rate, and work length), trials were first grouped by class. Within each neuron and class, trials were randomly partitioned into training (80%) and test (20%) pools. Pseudo-trials were then constructed independently for training and test sets by randomly drawing (with replacement) one trial from the corresponding pool for each neuron, and concatenating the resulting firing rates across neurons to form a single population vector. This procedure preserves class labels while approximating a simultaneously recorded population response. To equalize class evidence, we generated the same number of pseudo-trials per class (200 pseudo-trials per class, unless limited by available trials after splitting), and decoding performance was computed as the mean across classes.

Decoding was performed using a linear support vector machine (SVM) classifier trained on the pseudo-population vectors at each time bin. The entire pipeline (trial partitioning, pseudo-trial generation, model fitting, and evaluation) was repeated 100 times with independent random train/test splits to sample variability introduced by pseudo-population resampling. Reported decoding performance corresponds to the mean classification accuracy across iterations.

### Feedback geometry

#### Neural manifold visualization

To visualize the low-dimensional trajectory of population activity, neurons were first grouped by cortical area, and only those with sufficient trials in all feedback and speed conditions were included. For each area, condition-specific trial-averaged firing rates were computed and concatenated across neurons to form population matrices. We then apply the principal component analysis (PCA) to the z-scored population matrices, and only the first three principal components were used to visualize the temporal evolution of population activity across conditions.

#### Alignment of feedback encoding axes

For each bootstrap iteration and each rate *s*, we computed the population feedback axis *d*(*s*) = *v*_+_ (*s*)-*v*_−_ (*s*), where *v*_+_ (*s*) and *v*_−_ (*s*) are the pseudo-population mean activity vectors for positive and negative feedback at rate *s*, respectively (estimated from resampled pseudo-trials). We then computed cosine similarity between axes from different rates *cos*(*d*(*s*), *d*(*s*′)) and quantified rate-invariance of this axis by the mean off-diagonal cosine similarity *R*_*cos*_ = *mean*_*s*≠*s*′_ *cos*(*d*(*s*), *d*(*s*′)). Higher *R*_*cos*_ indicates that feedback axes are more parallel across rates.

#### Cross-condition decoding

We formed pseudo-populations by concatenating single-neuron activity across sessions. For each area (MCC, LPFC), trials were stratified by progress rate and feedback. Within a fixed analysis window, each neuron’s activity was averaged per trial. In each bootstrap, trials were split within each (rate × feedback) condition into train/test sets (80/20), and pseudo-trials were generated by sampling with replacement. A linear SVM was trained to classify feedback at one rate *s* and tested on each rate *s*′, yielding a 4×4 matrix *A* of accuracy. We summarized cross-rate generalization as 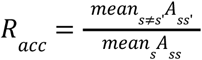, i.e., off-diagonal performance normalized by within-rate performance.

#### Null model (unit-identity shuffle across rates)

To test whether cross-rate generalization and axis alignment could arise without consistent cross-rate organization, we constructed a null by randomly permuting neuron identity independently for each rate. This procedure explicitly destroys the cross-rate neural manifold/alignment: it breaks any consistent mapping of dimensions (neurons) across rates, and therefore disrupts geometry that depends on neuron-wise correspondences. For decoding, we trained the classifier on observed data and evaluated it on test-rate pseudo-population vectors after applying the rate-specific unit permutation (mismatching feature identity across rates). For axis alignment, we applied the same per-rate unit permutations to the axes *d(s)* before computing *R*_*cos*_. One-sided p-values were computed as *p=*Pr(null≥observed).

#### MCC versus LPFC comparison

To compare areas fairly, we matched neuron counts by repeatedly subsampling the larger area down to the smaller area’s session count, recomputing *R*_*acc*_ and *R*_*cos*_ each time. We tested the one-sided hypothesis MCC > LPFC using the distribution of paired differences across subsamples.

### Timescale

To quantify each neuron’s intrinsic timescale (time constant, TAU), we followed the spike-autocorrelogram approach introduced by Fontanier et al. (2022)^24^, which estimates temporal signatures from continuous spike-time data rather than trial-epoch spike counts. This choice avoids the main limitation of the “classical” intrinsic-timescale method of Murray et al. (2014)^22^, which computes autocorrelation from binned spike counts within a fixed, repeated baseline epoch across trials.

#### Spike autocorrelogram construction

For each unit, we computed lagged differences between spike times up to the 100th successor spike (i.e., for each spike, the time differences to its next 1…100 spikes). These lagged differences were binned in 3.33 ms bins from 0–1000 ms to form an autocorrelogram, expressed as a probability density function. We removed the first 10 ms (to avoid refractory-period contamination) and smoothed the autocorrelogram using LOESS (span = 0.1). The autocorrelogram peak was defined as its maximum, except when the first bin was maximal, in which case the peak was taken as the first subsequent local maximum.

#### Exponential fit and TAU extraction

We fit the post-peak portion of each autocorrelogram with a mono-exponential decay plus offset, *AC*(*t*) = *Ae*^−*t*/τ^ + *B*, and took *τ* as the neuron’s intrinsic timescale (TAU). Fits were repeated 50 times with random initializations and the best fit was retained; only fits with positive *A, B*, and *τ* were accepted. For the small subset of units with a post-peak “dip” and secondary peak, Fontanier et al.’s conservative procedure was used to ensure the global exponential captured the final decay. Units without a valid global fit were excluded (MCC: 44/190 in monkey A, 25/104 in monkey H, LPFC: 84/168 in monkey A, 62/99 in monkey H).

#### Inclusion criterion

Because the autocorrelogram was estimated over a 0–1000 ms window and extreme estimates near the upper bound are less constrained, and consistent with the fact that frontal TAUs are typically reported within this range^22,24^, we restricted analyses to neurons with TAU < 900 ms.

#### Linking timescale to progress-rate encoding

For each neuron, we quantified progress-rate encoding strength as the absolute value of the slope coefficient from a univariate linear regression relating trial-by-trial firing rate to progress rate. For the time-resolved analysis, this quantity was computed separately at each time bin; for the window-based analysis, firing rates were first averaged within the predefined pre- and post-stimulus windows and the same regression was then applied. We next quantified, separately for each area and time window, the Pearson correlation between TAU and this encoding-strength measure across neurons. Significance was assessed with a permutation test (2000 permutations) in which the neuron-wise pairing between TAU and encoding strength was shuffled within monkey, preserving the monkey-level structure of the data, and permutation-based *p* values were reported together with the observed *r*.

### RNN model

We trained a recurrent neural network (RNN) to predict the internal gauge size on a trial-by-trial basis, using only information available to the animals at each trial. The model receives a sequential input stream corresponding to trials within a block and produces a prediction of the gauge size at each trial. Importantly, the true gauge size is only revealed on check trials, and supervision is applied only at those trials. We have chosen a number of hidden units to make a network that isn’t too large to outperform animals’ monitoring ability, nor too small to achieve the task nor allow lesioning of subsets of units. It furthermore allows us to compare decoding results as we have a similar amount of units.

At each trial *t*, the network receives the following inputs:

- Choice *c*_*t*_ ∈ {0, 1}, indicating the selected action.
- Feedback *f*_*t*_ ∈ {0, 1, 2}, encoding no feedback (check), correct work,incorrect work.
- Gauge size *g*_*t*_ ∈ {1,…, 7}, which equals the true gauge size on check trials and is set to a null symbol on work trials.

Thus, although a gauge input channel is always present, it carries informative values only on check trials, mimicking the experimental task structure. The network must therefore infer the gauge evolution internally across work trials. Each discrete input is then embedded using a learned linear embedding.

The network core consists of a single-layer LSTM with hidden state *h*_*t*_ and cell state *c*_*t*_ :

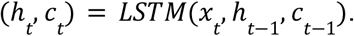

At each trial, the network predicts a categorical distribution over gauge sizes *k* ∈ {1,…, 7}:

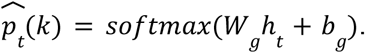

The predicted gauge size is defined as the maximum a posteriori estimate:

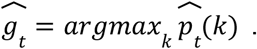

Crucially, this prediction is computed before any gauge observation at trial *t*, ensuring causal prediction.

The network is trained using negative log-likelihood loss applied only on check trials:

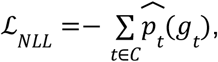

where *C* denotes the set of check trials. To encourage stable internal dynamics, we add a smoothness regularization term on the hidden state:

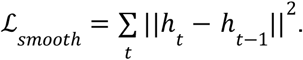

In addition, to enforce the task contingency that incorrect work trials should not advance progress, we include a penalty that discourages positive gauge updates following incorrect feedback without hard-coding the update rule:

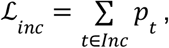

where *p*_*t*_ denotes the network’s predicted gauge increment (or increment propensity) at trial *t*, and *Inc* denotes the set of incorrect work trials.

The total loss is a weighted sum: ℒ = ℒ_*NLL*_ + *λ*_*smooth*_ ℒ_*smooth*_ + *λ*_*inc*_ ℒ_*inc*_. Training is performed with AdamW optimization, gradient clipping, and early stopping based on validation loss.

#### Progress-rate decoding

To assess whether progress rate information is represented in the hidden state, we performed linear decoding of the block’s progress rate from *h*_*t*_. Hidden states were extracted before any check, as well as before successive checks within a block. For each check index *k*, we trained a multinomial logistic regression classifier to predict the progress rate (labels 1–4), using stratified cross-validation. To ensure fair comparisons across conditions (lesion/no lesion), decoding analyses were performed on the same blocks and same cross-validation folds.

#### Lesion analysis

Progress-rate units were identified by regressing hidden states onto the block-wise progress rate. Units were ranked by the absolute value of their regression weights aggregated across trials. Lesions were implemented by silencing selected hidden units: 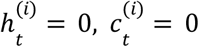 for all trials *t*, where *i* indexes lesioned units.

To force the network to rely on its internal belief instead of gauge size input at check trials to predict the gauge size, we evaluated gauge prediction error under a masked-observation regime: After the second check, the gauge observation channel was masked. Performance was quantified using mean squared error (MSE) across all trials, averaged across blocks.

Statistical significance was assessed using a one-sided permutation test:

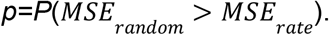

#### Feedback modulation

To characterize how the network integrates feedback, we measured the expected change in predicted gauge size following work trials. This quantity was computed separately for correct and incorrect work trials and stratified by progress rate. This analysis reveals whether the magnitude of belief updates depends jointly on feedback valence and progress rate, analogous to feedback-modulated neural responses observed experimentally.

Finally, we examined whether feedback representations generalize across progress rates. For each rate, a linear decoder was trained to classify feedback (correct vs incorrect) from hidden states. Decoders trained at one rate were tested on data from other rates, yielding a 4×4 cross-rate generalization matrix. Perfect cross-rate generalization indicates a shared, rate-invariant feedback axis in the hidden state space, while deviations indicate rate-dependent feedback encoding.

---

**Extended Figure 1.**
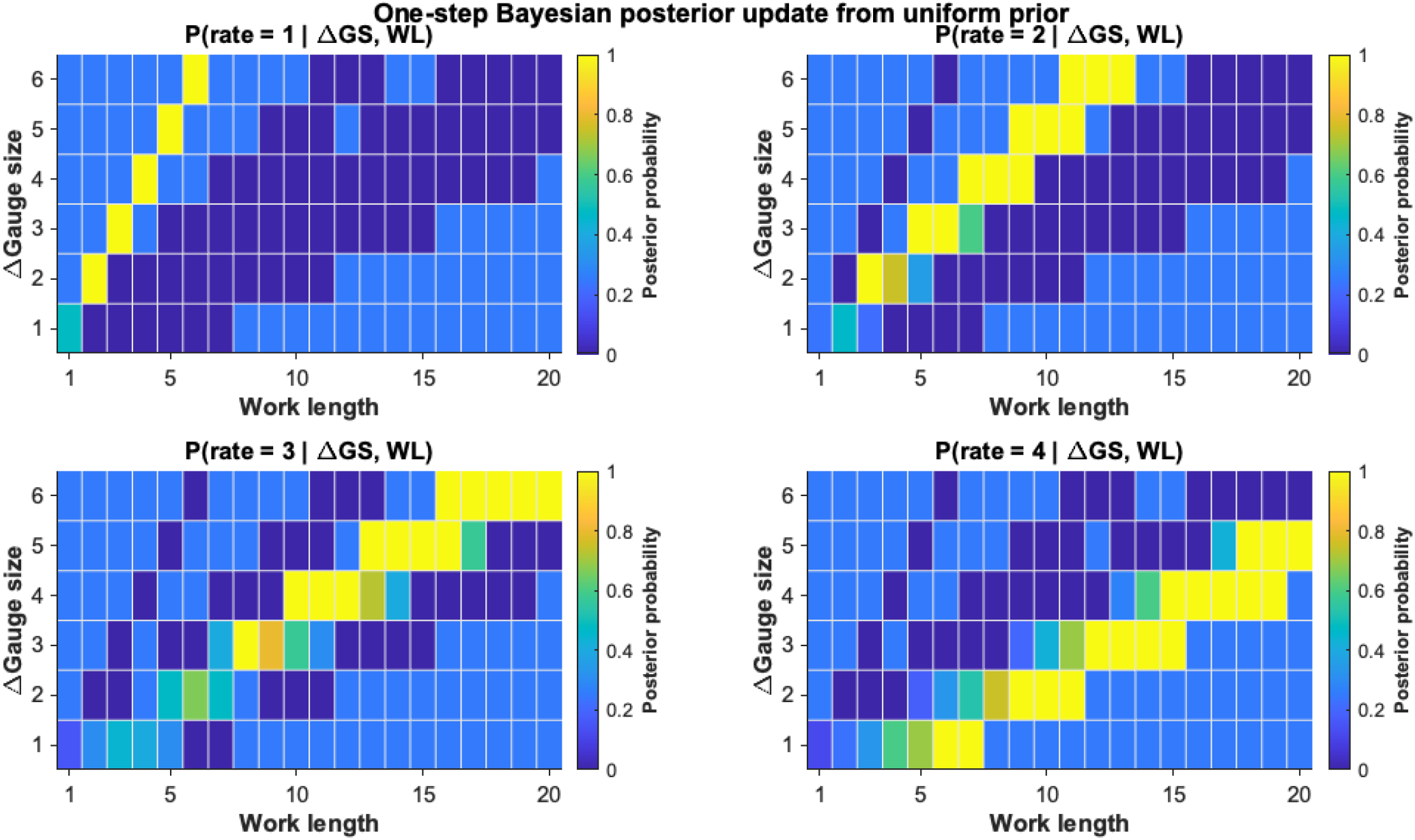
The posterior probability of progress rate (indicated by the colour) given the observed change of gauge size between checks (row) and the work length (column).

**Extended Figure 2.**
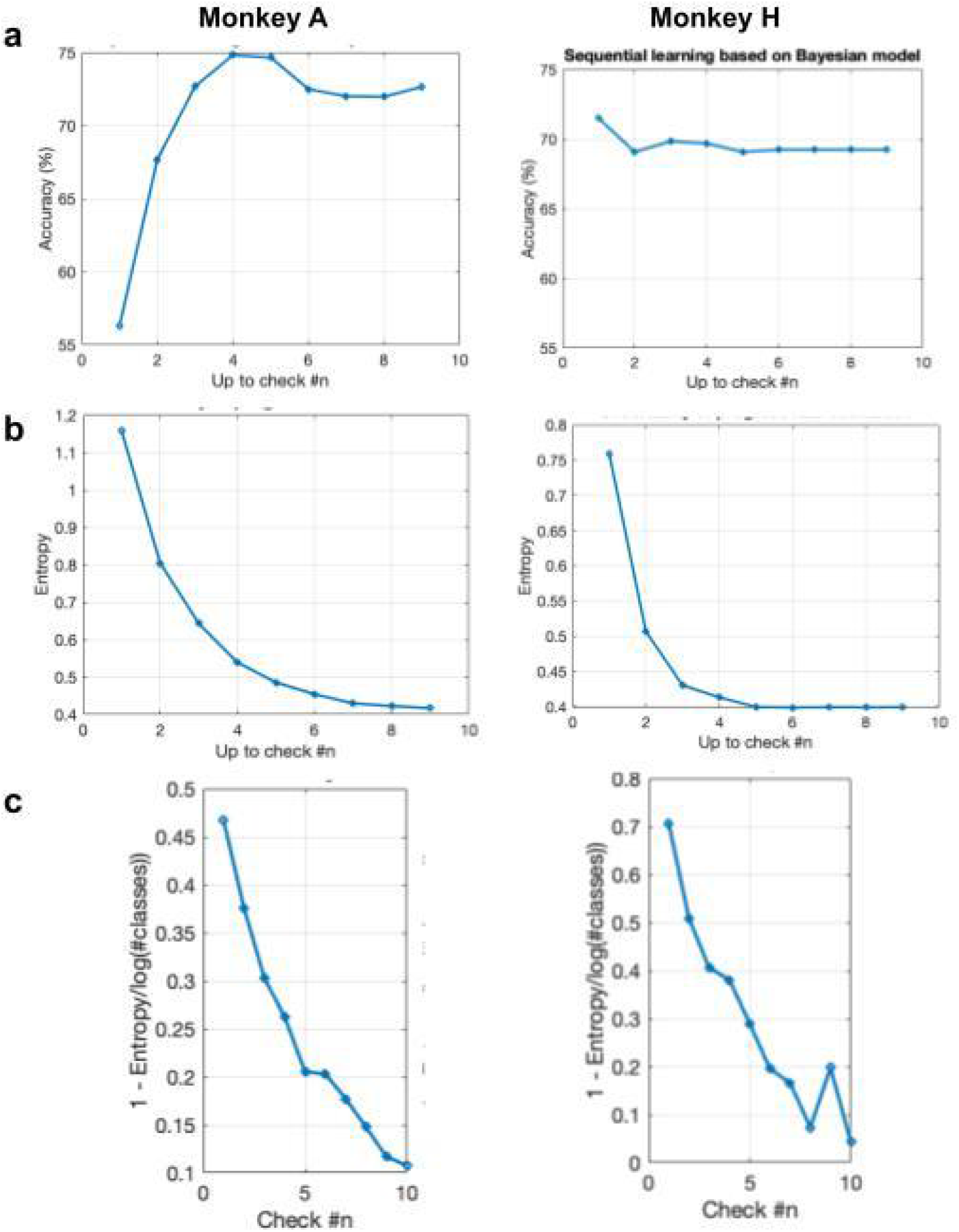
(a) Bayesian progress rate estimation accuracy compared to ground truth as a function of the number of checks. (b) Entropy of the posterior distribution over progress rates as a function of the number of checks. (c) The contribution of each check to reducing uncertainty in progress rate estimation, measured by the decrease in entropy.

**Extended Data Figure 3.**
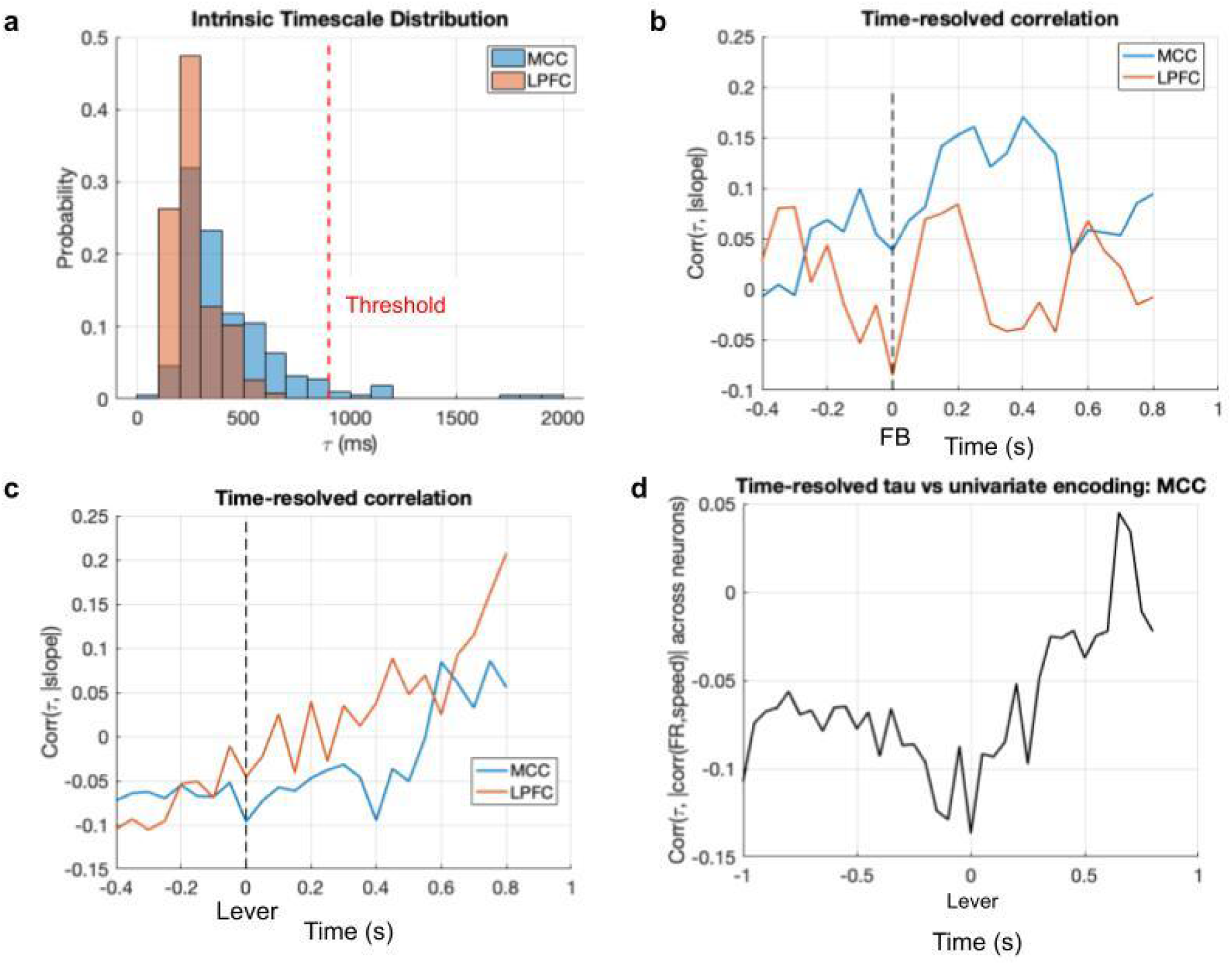
(a) Distributions of intrinsic neuronal timescales ***τ*** estimated from spike-count autocorrelation for MCC and LPFC (see methods). The red dashed line indicates the exclusion threshold used in the following correlation analysis. (b) Time-resolved correlation between each neuron’s intrinsic timescale and the magnitude of its progress-rate encoding (regression slope magnitude), aligned to feedback onset. (c) Time-resolved correlation between each neuron’s intrinsic timescale and the magnitude of its progress-rate encoding (regression slope magnitude), aligned to lever onset. (d) same as (c) but only using check trials and MCC neurons.

